# Proteomic signatures of *APOE* ε4 across human tissues and cell types in Alzheimer’s disease

**DOI:** 10.64898/2026.08.27.746928

**Authors:** Xinran C. Li, Caitlin A. Finney, Artur Shvetcov

**Author notes:** Corresponding authors: Email.; Address. 176 Hawkesbury Road, Westmead, New South Wales, Australia 2145. Equal senior authors (Caitlin A. Finney, Artur Shvetcov).

## Abstract

The apolipoprotein E ε4 (*APOE* ε4) allele is the strongest genetic risk factor for late-onset Alzheimer’s disease (AD), yet its associated protein changes across biofluids, brain regions, and cell types remain incompletely understood. This study included 1691 participants from the Religious Orders Study and Rush Memory and Aging Project (ROSMAP), 1226 participants from the Accelerating Medicines Partnership - Alzheimer’s Disease (AMP-AD) Diverse Cohorts Study, and 735 participants from the Alzheimer’s Disease Neuroimaging Initiative (ADNI). To characterise *APOE* ε4 effects, we analysed proteomic data from plasma, cerebrospinal fluid (CSF), and patient induced pluripotent stem cell (iPSC)-derived astrocytes and neurons, as well as transcriptomic and proteomic data from multiple brain regions. In plasma, *APOE* ε4 carriers shared a proteomic signature enriched for immune processes, irrespective of AD diagnosis. A machine learning classifier trained on this plasma signature discriminated carriers from non-carriers in an independent CSF cohort, and mediation analysis resolved a subset of these proteins into those lying upstream and downstream of AD. Carriage was also associated with greater neuropathological burden, reflected in higher Braak stages and CERAD scores. However, only limited *APOE* ε4-associated transcriptomic and proteomic changes were observed in bulk brain tissue, with poor concordance between the transcriptome and proteome. Proteomic analyses of iPSC-derived astrocytes and neurons further revealed cell-type-specific *APOE* ε4-associated changes. These findings show that *APOE* ε4 is associated with a consistent proteomic signature across plasma and CSF, but its effects in the brain differ across cell types and brain regions.

## Background

The apolipoprotein E (*APOE*) ε4 allele is the strongest genetic risk factor for late-onset Alzheimer’s disease (AD), increasing disease risk in a dose-dependent manner and lowering the age at onset (1, 2). *APOE* ε4 carriage is also associated with earlier and more severe accumulation of amyloid plaques and neurofibrillary tangles (2, 3, 4). Despite this well-established link, the molecular mechanisms through which *APOE* ε4 contributes to AD pathogenesis remain incompletely understood, limiting opportunities for targeted preventative or therapeutic strategies in *APOE* ε4 carriers.

Recent advances in high-throughput transcriptomics and proteomics have offered important insights into *APOE* ε4-associated molecular mechanisms that may contribute to AD. In plasma and cerebrospinal fluid (CSF), proteomic studies have consistently reported *APOE* ε4-associated changes implicating immune-related dysregulation (5, 6, 7, 8, 9, 10). *APOE* ε4-associated effects have also been reported in the brain. Transcriptomic studies have identified mRNA changes related to immune, metabolic, and cerebrovascular function, with effects varying across cell types and brain regions examined (11, 12, 13, 14, 15). Similarly, proteomic studies have identified changes associated with synaptic, metabolic or immune function depending on the brain regions and cohorts examined (5, 9, 16).

Although studies report a moderate correlation between *APOE* ε4-associated protein changes across plasma and CSF (17), whether the same protein signature is preserved across biofluids remains unknown. Resolving this would clarify whether plasma, which is more accessible than CSF, can capture *APOE* ε4-associated changes relevant to the central nervous system. It is also unclear whether *APOE* ε4-associated plasma proteins lie upstream of AD, where they may contribute to disease risk, or arise as downstream consequences of established pathology. In the brain, *APOE* ε4 is a well-established driver of amyloid and tau pathology (2, 3, 4, 18). Bulk transcriptomic and proteomic studies, however, have reported comparatively modest *APOE* ε4-associated changes (19). Whether this reflects a genuine absence of effect or the masking of cell type-specific changes within heterogeneous bulk tissue remains unknown. These findings are further complicated by limited mRNA-protein concordance in the human brain (20, 21, 22, 23). The relationship between the brain transcriptome and proteome has not been examined in relation to *APOE* ε4, making it unclear the extent to which *APOE* ε4-associated transcriptomic changes are reflected at the protein level.

Here, we address these gaps through a comprehensive characterisation of *APOE* ε4-associated molecular changes across plasma, CSF, post-mortem brain tissue, and induced pluripotent stem cell (iPSC)-derived neurons and astrocytes from three independent AD cohorts (Fig. 1). We first develop a machine learning classifier using *APOE* ε4-associated plasma proteins and show that it generalises to CSF. Mediation analysis further reveals a subset of *APOE* ε4-associated plasma proteins that may interact with AD. Next, we show that *APOE* ε4 carriers with cognitive impairment show greater neuropathological burden in the brain. In contrast, we observe limited *APOE* ε4-associated transcriptomic and proteomic changes in bulk tissue across multiple brain regions. We further find that concordance between *APOE* ε4-associated mRNA and protein changes is limited, highlighting that protein abundance cannot be reliably inferred from mRNA expression. Finally, we offer a potential explanation for the limited molecular changes observed in bulk brain tissue by identifying cell type-specific proteomic signatures in iPSC-derived astrocytes and neurons from the same cohorts. These findings suggest that bulk tissue profiling may dilute cell type-specific changes. Together, this study provides a comprehensive view of the molecular landscape associated with *APOE* ε4 carriage across tissues and cell types.

**Figure 1.**
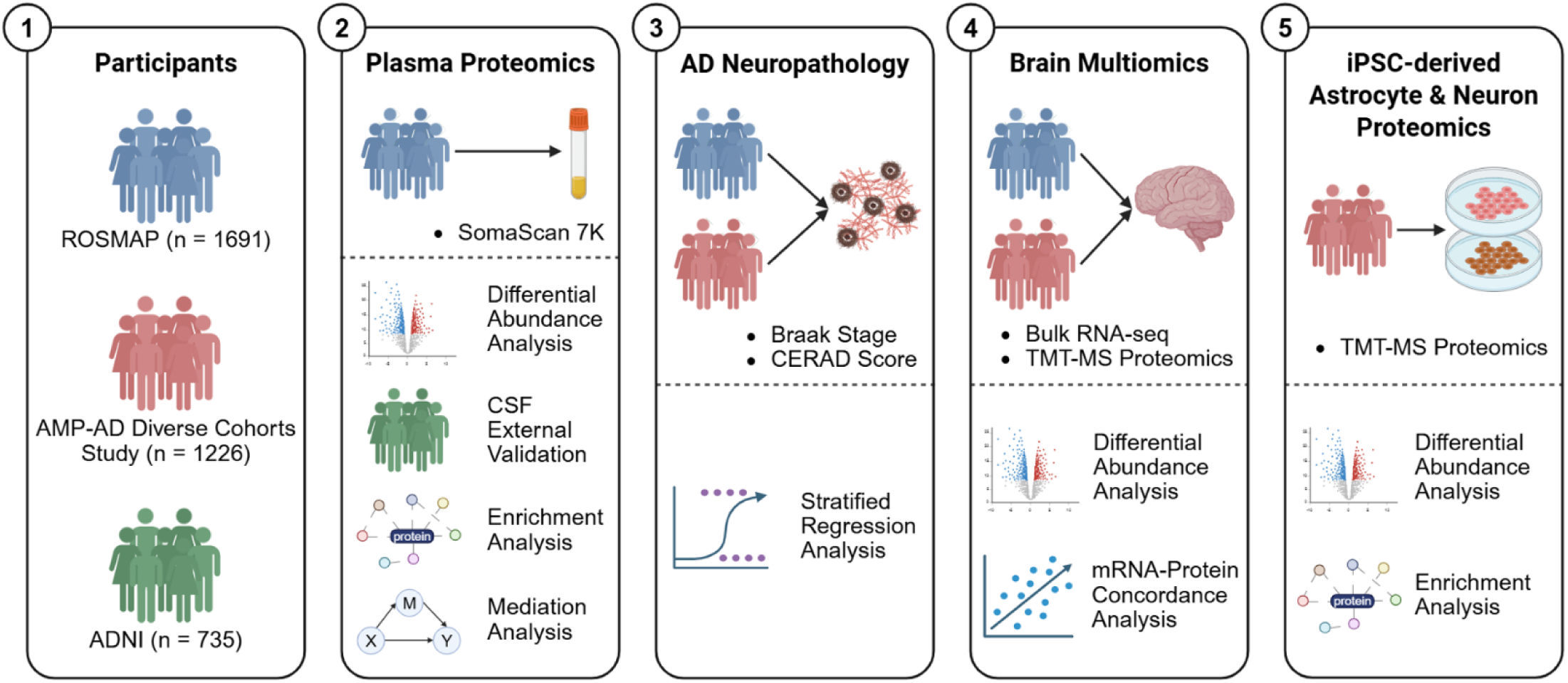
Overview of the study design. Data from ROSMAP, AMP-AD Diverse Cohorts Study, and ADNI were sourced. Plasma and CSF proteomics, AD neuropathology, brain transcriptomics and proteomics, and proteomics from iPSC-derived astrocytes and neurons were analysed. AD, Alzheimer’s disease; ADNI, Alzheimer’s Disease Neuroimaging Initiative; AMP-AD, Accelerating Medicines Partnership for Alzheimer’s Disease; iPSC, induced pluripotent stem cell; ROSMAP, Religious Orders Study and Memory and Aging Project.

## Methods Participants

### ROSMAP

The Religious Orders Study (ROS) and Rush Memory and Aging Project (MAP) are two longitudinal cohort studies of ageing and AD conducted by the Rush Alzheimer’s Disease Centre (24). ROS, initiated in 1994, enrolled older Catholic clergy from across the United States, and MAP, initiated in 1997, enrolled community-dwelling older adults from northeastern Illinois. All participants underwent annual clinical and cognitive assessments and provided informed consent to post-mortem brain donation, as described elsewhere (24). The present study utilised proteomic, transcriptomic, and neuropathological data from 1691 participants (69% female, 25% carrying one or more *APOE* ε4 alleles). Among them, 697 had no cognitive impairment (NCI), 380 had mild cognitive impairment (MCI), and 614 had AD at their last available assessment. Diagnoses were made via a three-stage process comprising cognitive testing, neuropsychological evaluation, and clinician judgement (25). For plasma proteomic analyses, diagnoses at the time of plasma collection were used, and when participants had multiple visits, only the earliest available measurement was retained. For all other analyses, diagnosis at death was used. Demographic characteristics of each analytic subset are provided in Supplementary Table 1.

#### AMP-AD Diverse Cohorts Study

The Accelerating Medicines Partnership - Alzheimer’s Disease (AMP-AD) Diverse Cohorts Study is a cross-consortium project generating harmonised post-mortem multi-omic data from ancestrally diverse human cohorts (26). The present study utilised proteomic, transcriptomic, and neuropathological data from 1226 participants (60% female, 36% carrying one or more *APOE* ε4 alleles). Among them, 389 had NCI and 837 had AD. Diagnoses were confirmed post-mortem via neuropathological evaluation, as described elsewhere (27). Demographic characteristics of each analytic subset are provided in Supplementary Table 2.

#### ADNI Study

The Alzheimer’s Disease Neuroimaging Initiative (ADNI) is a multisite longitudinal study tracking progression from normal cognition to AD using neuroimaging and biomarkers (28). In this study, SomaLogic CSF proteomics data from 735 ADNI participants were used as an independent external validation cohort for the machine learning classifier.

The sample comprised 143 participants with NCI, 283 with MCI, and 309 with AD, of whom 49% carried one or more *APOE ε4* alleles. Diagnoses were made based on Mini-Mental State Examination (MMSE) and Clinical Dementia Rating (CDR) scores, as described elsewhere (28). Detailed demographic characteristics are provided in Supplementary Table 3.

#### Neuropathology

Neurofibrillary tangle burden and neuritic plaque burden were quantified by Braak stage and Consortium to Establish a Registry for Alzheimer’s Disease (CERAD) score, respectively, as described previously (29, 30). Associations between *APOE* ε4 carrier status and each neuropathological burden were assessed separately within each diagnostic group using ordinal or binary logistic regression, adjusted for age at death and sex. In the ROSMAP cohort, Braak stage was grouped into three ordinal categories prior to analysis (None - Stage II, Stages III-IV, and Stages V-VI) to ensure sufficient cell counts within each category. In the AMP-AD Diverse Cohorts Study, categories with no observations within a given diagnostic group were excluded prior to model fitting. Results are reported as odds ratios (ORs) with 95% confidence intervals (CIs). Analyses were implemented using the ‘MASS’ package in R (v4.5.2) (31).

### Transcriptomic and Proteomic Data Preprocessing

#### Bulk brain tissue transcriptomics

Bulk RNA-sequencing (RNA-seq) data from the ROSMAP cohort were generated from dorsolateral prefrontal cortex (dlPFC), head of caudate nucleus (hCN), and posterior cingulate cortex (PCC) using the Illumina HiSeq 2000 platform, as previously described (32, 33). Bulk RNA-seq data from the AMP-AD Diverse Cohorts Study were generated from the dlPFC and superior temporal gyrus (STG). Samples were sourced from four contributing sites (Mayo-Emory, Columbia, MSSM, and Rush), each sequenced using the Illumina NovaSeq 6000 platform as previously described (27). Data were merged across sites, retaining only genes present across all four contributing sites for downstream analysis. For quality control, genes were excluded if they had fewer than 10 counts in at least *n* samples, where *n* was the number of samples in the smaller of the two *APOE* ε4 groups.

#### Plasma and CSF proteomics

Plasma proteomic data from the ROSMAP cohort and CSF proteomic data from the ADNI cohort were both generated using the SomaScan 7K aptamer-based platform (SomaLogic) as previously described (24, 34). All samples were pre-processed using SomaLogic’s standard multi-step normalisation pipeline, including adaptive normalisation by maximum likelihood (ANML), to reduce technical variability (35). Protein abundances were quantified as relative fluorescence units (RFU) and log10-transformed prior to downstream analysis. Each aptamer was mapped to a UniProt identifier, and aptamers that did not map to a UniProt identifier or had duplicate mappings were excluded.

#### Bulk brain tissue proteomics

dlPFC proteomic data from the ROSMAP cohort and AMP-AD Diverse Cohorts Study were used. All datasets were generated using tandem mass tag mass spectrometry (TMT-MS) and normalised as previously described (27, 36, 37). For all datasets, proteins with more than 30% missing values were excluded, and remaining missing values were imputed using the within-protein median. Protein abundances were log2-transformed prior to downstream analysis.

#### iPSC-derived astrocytes and neurons proteomics

iPSC lines were generated from cryopreserved peripheral blood mononuclear cells (PBMCs) collected from ROSMAP participants as previously described (38). iPSC lines were differentiated into iAstrocytes via lentiviral co-overexpression of SOX9 and NFIB and profiled using TMT-MS, as previously described (39). iPSC lines were differentiated into iNeurons via lentiviral overexpression of NGN2 and profiled using TMT-MS, as previously described (38). For iAstrocyte proteomics, protein abundances were provided in normalised, log2-transformed values with no missing data. Batch-associated variation was removed using the removeBatchEffect function in the ‘limma’ package in R, with sex and diagnosis included as covariates in the design matrix (40). For iNeuron proteomics, proteins with more than 30% missing values were excluded, and remaining missing values were imputed using the within-protein median. Protein abundances were then log2-transformed. Each sample was median-centred to correct for between-sample differences in total protein abundance prior to downstream analysis.

## Statistical Analyses

### Mutual information-based feature selection

To identify *APOE* ε4-associated proteins, mutual information (MI) was calculated between protein abundance and *APOE* ε4 carriage. This captures both linear and non-linear relationships, with higher MI values indicating a stronger statistical association (9, 41). For plasma proteomics, MI was computed exclusively in NCI participants to minimise confounding by AD pathology, and proteins exceeding an MI threshold of 0.1 were selected to train machine learning classifiers (9). For brain proteomics and transcriptomics, MI was similarly computed in NCI participants. For iAstrocyte and iNeuron proteomics, where sample sizes were smaller, MI was computed across all available samples. The 13 proteins with the highest non-zero MI scores were selected per cell type, and principal component analysis (PCA) was subsequently performed to evaluate whether MI-selected proteins separated participants by *APOE* ε4 carriage. All MI calculations were performed in R (v4.5.2) using the ‘FSelectorRcpp’ package.

### Random forest-based machine learning classifier

To evaluate whether the *APOE* ε4-associated proteomic signature identified in NCI participants was preserved across diagnostic groups, random forest classifiers were trained on NCI participants using five-fold cross-validation repeated five times, with class imbalance addressed using the Synthetic Minority Oversampling Technique (SMOTE) (42). Predictive performance was assessed in MCI and AD cohorts using sensitivity, specificity, positive predictive value (PPV), negative predictive value (NPV), and area under the receiver operating characteristic curve (AUC). To examine whether the signature was confounded by sex, a separate classifier using the same proteins was trained exclusively on female participants and tested in male participants. To assess generalisation beyond plasma and the ROSMAP cohort, a classifier was trained on the full ROSMAP sample and tested on an independent CSF proteomic dataset from ADNI. All classifier training and testing were performed using the ‘caret’ package in R (43).

### Differential abundance analysis

Differentially abundant proteins (DAPs) were identified by fitting empirical Bayes-moderated linear models using the ‘limma’ package in R, with adjustment for age, sex, diagnosis, and batch where applicable (40). Differentially expressed genes (DEGs) were identified by fitting negative binomial generalised linear models using the ‘DESeq2’ package in R, with adjustment for age, sex, diagnosis, batch and data-contributing site (44). For both analyses, statistical significance was determined using a false discovery rate (FDR)-adjusted *p*-value below 0.05, calculated using the Benjamini-Hochberg procedure.

### mRNA-protein concordance

To assess concordance between mRNA and protein changes in the dlPFC, we first identified participants in the AMP-AD Diverse Cohorts Study with both transcriptomic and proteomic data available. Ensembl gene IDs and UniProt protein IDs were mapped to a common gene symbol space using the ‘biomaRt’ package in R (45). For all mRNA-protein pairs, Spearman correlation coefficients were computed between transcriptomic and proteomic log2 fold changes associated with *APOE* ε4 carriage and AD status, separately.

### Mediation analysis

To assess the directional relationship between *APOE* ε4-associated plasma proteins and AD pathology, mediation analyses were performed in participants with NCI or AD using the ‘mediation’ package in R (46). Candidate plasma proteins were first identified through differential abundance analysis adjusted for age and sex, using an FDR threshold of 0.05. Two causal pathways were evaluated for each candidate protein: an upstream model in which the protein mediates the effect of *APOE* ε4 on AD status (*APOE* ε4 ◊ protein ◊AD), and a downstream model in which AD status mediates the effect of *APOE* ε4 on protein abundance (*APOE* ε4 ◊ AD ◊ protein). In the upstream model, linear regression was used for the mediator (protein abundance) and logistic regression for the outcome (AD status). In the downstream model, logistic regression was used for the mediator (AD status) and linear regression for the outcome (protein abundance). All regressions were adjusted for age at visit and sex. Mediation effects with 95% CIs were estimated using 1000 bootstrap resamples and resulting *p*-values were FDR-adjusted for multiple testing.

### Functional enrichment analysis

Functional enrichment analyses were performed using NetworkAnalyst (v3.0) with all proteins passing preprocessing used as the background universe (47, 48, 49). Protein-protein interaction networks were constructed using a first-order network in the International Molecular Exchange (IMEx) Consortium Interactome, followed by minimum network simplification (50). Enrichment of biological processes was assessed using the Gene Ontology (GO) Biological Process database, and pathway-level enrichment was assessed using the Reactome database. Statistical significance was defined as an FDR-adjusted *p*-value below 0.05.

## Results

### *APOE* ε4 carriers share a distinct plasma proteomic signature independent of AD pathology

We first sought to characterise plasma proteomic signatures associated with *APOE* ε4 carriage (Fig. 1). Linear models adjusted for age, sex, and diagnosis identified 53 *APOE* ε4-associated DAPs among 793 ROSMAP participants with plasma proteomic data (Fig. 2a; Supplementary Table 4). PCA visualisation restricted to these 53 DAPs revealed clear separation by *APOE* ε4 carriage compared with PCA using all 6402 proteins (Fig. 2b-c).

**Figure 2.**
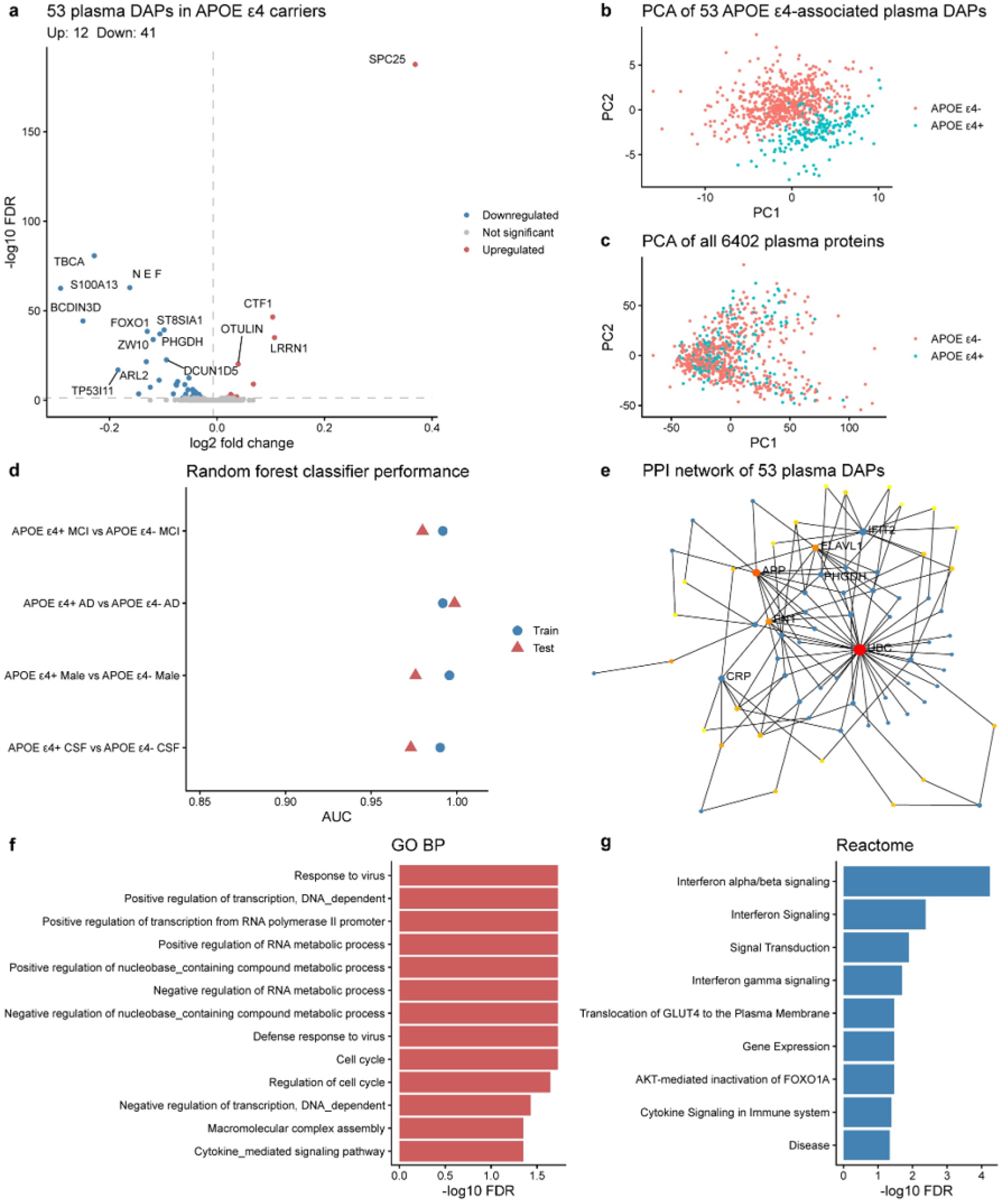
*APOE* ε4-associated plasma proteomic signature in ROSMAP participants. (a) Volcano plot of 53 *APOE* ε4-associated DAPs identified across all ROSMAP participants with plasma proteomics (FDR < 0.05). (b) PCA of the 53 APOE ε4-associated DAPs shows clear separation by *APOE* ε4 carriage. (c) PCA of all 6402 quantified proteins shows no clear separation by *APOE* ε4 carriage. (d) AUC dot plot showing classifier performance across training and test sets for four generalisation scenarios. (e) Protein-protein interaction network of the 53 *APOE* ε4-associated DAPs; blue nodes represent identified DAPs, and other coloured nodes represent imputed interacting proteins. (f) GO biological process enrichment results (FDR < 0.05). (g) Reactome pathway enrichment results (FDR < 0.05). AMP-AD, Accelerating Medicines Partnership - Alzheimer’s Disease; APOE, apolipoprotein E; AUC, area under the receiver operating characteristic curve; DAP, differentially abundant protein; FDR, false discovery rate; GO BP, gene ontology biological processes; PCA, principal component analysis; ROSMAP, Religious Orders Study and Memory and Aging Project.

We next used machine learning to examine whether this *APOE* ε4-associated proteomic signature was independent of cognitive impairment. To identify a parsimonious protein set for model training that was not confounded by AD pathology, MI-based feature selection was applied to data from NCI participants (*n* = 420). This identified 10 proteins with a MI exceeding 0.1: SPC25, TBCA, NEFL, LRRN1, S100A13, ST8SIA1, BCDIN3D, FOXO1, PHGDH, and ZW10 (Supplementary Table 5). All 10 proteins were among the 53 DAPs identified earlier. A random forest classifier trained on NCI participants using these 10 proteins reliably discriminated *APOE* ε4 carriers from non-carriers in participants with MCI (AUC = 0.98) and AD (AUC = 1.00), suggesting this signature was independent of cognitive impairment (Table 1; Fig. 2d). To determine whether the signature was influenced by sex, a classifier trained on female participants was tested in male participants and retained strong discriminative performance (AUC = 0.98) (Table 1; Fig. 2d). To assess whether the signature generalised beyond plasma and the ROSMAP cohort, we tested the classifier on an independent CSF proteomic dataset from ADNI. This achieved comparably strong performance (AUC = 0.97) (Table 1; Fig. 2d).

**Table 1.** Performance metrics of random forest classifiers evaluating the generalisability of the *APOE* ε4-associated proteomic signature identified in individuals with no cognitive impairment.

| Comparison | Sensitivity | Specificity | PPV | NPV | AUC |
| --- | --- | --- | --- | --- | --- |
| *APOE* $\epsilon 4+$ MCI vs *APOE* $\epsilon 4-$ MCI | 0.92 | 0.99 | 0.97 | 0.97 | 0.98 |
| *APOE* $\epsilon 4+$ AD vs *APOE* $\epsilon 4-$ AD | 0.90 | 1.00 | 1.00 | 0.94 | 1.00 |
| *APOE* $\epsilon 4+$ Male vs *APOE* $\epsilon 4-$ Male | 0.95 | 0.96 | 0.90 | 0.98 | 0.98 |
| *APOE* $\epsilon 4+$ CSF vs *APOE* $\epsilon 4-$ CSF | 0.96 | 0.87 | 0.88 | 0.96 | 0.97 |

Abbreviations: AD, Alzheimer's Disease; APOE, apolipoprotein E; AUC, area under the receiver operating characteristic curve; CSF, cerebrospinal fluid; MCI, mild cognitive impairment; NPV, negative predictive value; PPV, positive predictive value.

We then characterised the biological function of *APOE* ε4-associated DAPs using a protein-protein interaction network analysis. This revealed functional connectivity among the 53 proteins (Fig. 2e). GO biological processes enrichment analysis identified significant (FDR < 0.05) overrepresentation of processes related to inflammatory responses, transcriptional regulation, RNA metabolic processes, and cell cycle regulation (Fig. 2f). Reactome pathway enrichment analysis further revealed strong overrepresentation of interferon signalling and other pathways related to the immune system (Fig. 2g).

Together, these findings indicate that *APOE* ε4 carriers share a distinct plasma proteomic signature that is robust across diagnostic groups and sex, and generalises to CSF proteomics from an independent cohort. These *APOE* ε4-associated proteins are functionally interconnected and have strong implications in inflammatory and immune-related processes.

### Mediation analysis reveals bidirectional relationships between *APOE* ε4-associated plasma proteins and AD pathology

We next sought to investigate whether *APOE* ε4-associated plasma proteins act as upstream mediators of AD risk or reflect downstream consequences of AD pathology (Fig. 3a). Using linear models adjusted for age and sex, we first identified 43 *APOE* ε4-associated DAPs in ROSMAP participants with NCI or AD (Fig. 3b). Each protein was evaluated in two mediation pathways: *APOE* ε4 ◊ protein ◊ AD and *APOE* ε4 ◊ AD ◊ protein (Fig. 3a). Of these, 10 proteins showed a significant average causal mediation effect (ACME) in at least one pathway after FDR correction (Fig. 3c; Supplementary Tables 6-7). CTF1 showed complementary mediation in both pathways, consistent with a bidirectional statistical association in which *APOE* ε4-associated upregulation of CTF1 may contribute to AD risk, and AD pathology further elevates CTF1 abundance. Six proteins (NEFL, BCDIN3D, ST8SIA1, DCUN1D5, TP53I11, and KCTD2) showed competitive mediation in both pathways. *APOE* ε4 was associated with lower levels of these proteins, and lower levels were associated with lower AD risk. In the reverse direction, AD pathology was associated with higher levels of the same proteins, partially offsetting the *APOE* ε4-associated reduction. IFIT2 showed competitive mediation only in the *APOE* ε4 ◊ protein ◊ AD pathway, consistent with a role in attenuating the effect of *APOE* ε4 on AD. CDA and BIRC2, however, showed significant mediation only in the *APOE* ε4 ◊ AD ◊ protein pathway, indicating that their altered abundance in *APOE* ε4 carriers is partly a downstream consequence of AD pathology. Together, these findings reveal a subset of *APOE* ε4-associated plasma proteins that may statistically mediate, reflect, or bidirectionally associate with AD, while the majority showed no significant association with AD in either direction.

**Figure 3.**
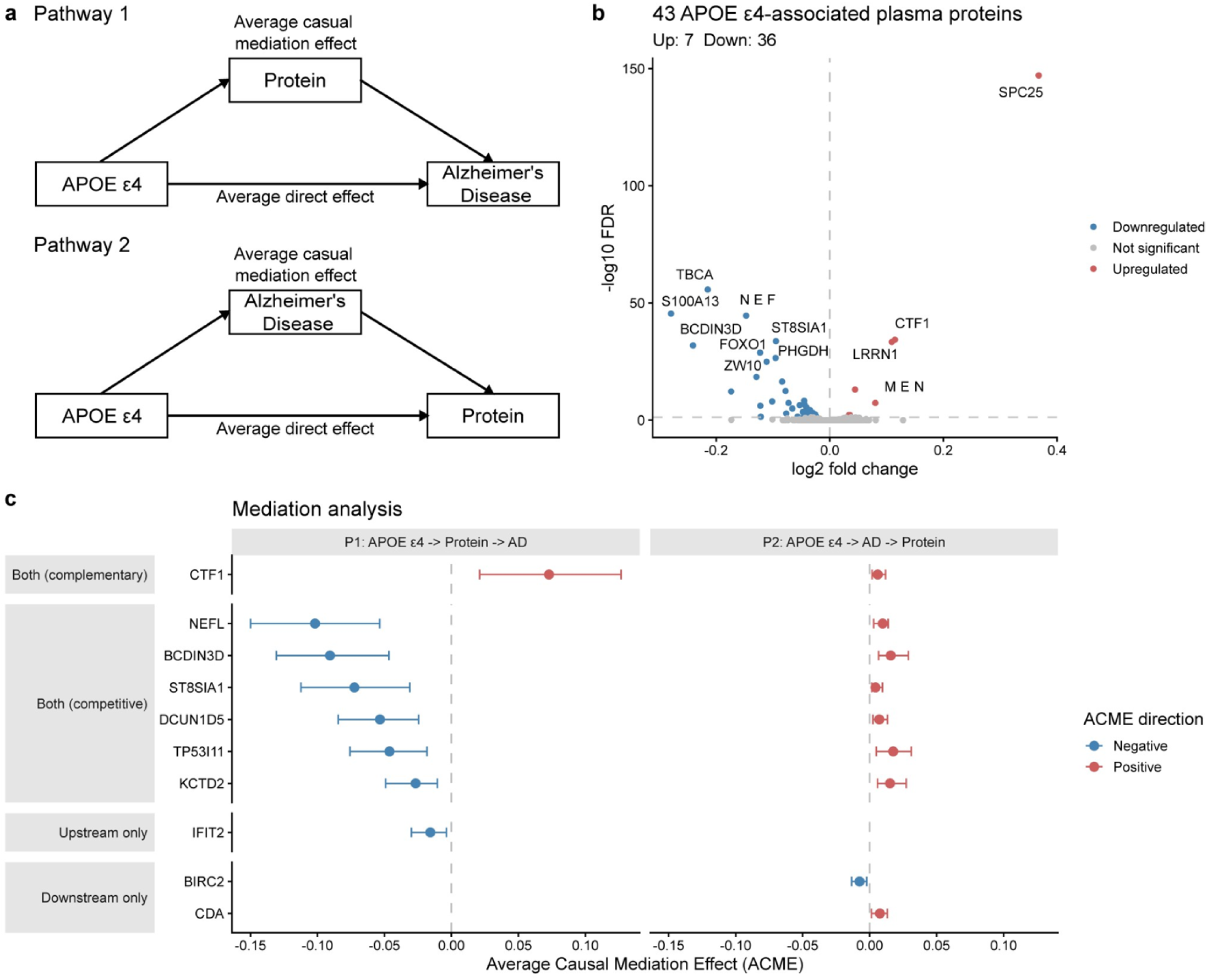
Mediation analysis reveals relationships between *APOE* ε4-associated plasma proteins and Alzheimer’s disease. (a) Schematic of the two mediation pathways tested. (b) Volcano plot of 43 *APOE* ε4-associated DAPs identified in ROSMAP participants with NCI or AD, without adjustment for AD diagnosis (FDR < 0.05). (c) Forest plot of ACME estimates and 95% bootstrap confidence intervals for the 10 proteins with a significant ACME in at least one pathway (FDR < 0.05). ACME, average causal mediation effect; AD, Alzheimer’s disease; APOE, apolipoprotein E; DAP, differentially abundant protein; FDR, false discovery rate; NCI, no cognitive impairment; ROSMAP, Religious Orders Study and Memory and Aging Project.

### *APOE* ε4 carriage is associated with greater AD neuropathological burden in individuals with cognitive impairment

To investigate whether *APOE* ε4 effects extend to brain pathology, we examined the association between *APOE* ε4 carriage and neurofibrillary tangle burden (Braak stage) and neuritic plaque burden (CERAD score) across diagnostic groups. In the ROSMAP cohort, ordinal regression adjusted for age and sex showed that *APOE* ε4 carriage was significantly associated with higher Braak stage in the MCI (OR = 2.36, 95% CI: 1.36 - 4.11) and AD groups (OR = 2.58, 95% CI: 1.82 - 3.68), but not in the NCI group (OR = 1.10, 95% CI: 0.66 - 1.83) (Fig. 4a). In contrast, APOE ε4 carriage was associated with more severe CERAD scores across all three diagnostic groups (NCI: OR = 2.71, 95% CI: 1.76 - 4.20; MCI: OR = 2.71, 95% CI: 1.65 - 4.50; AD: OR = 3.37, 95% CI: 2.36 - 4.87) (Fig. 4b).

**Figure 4.**
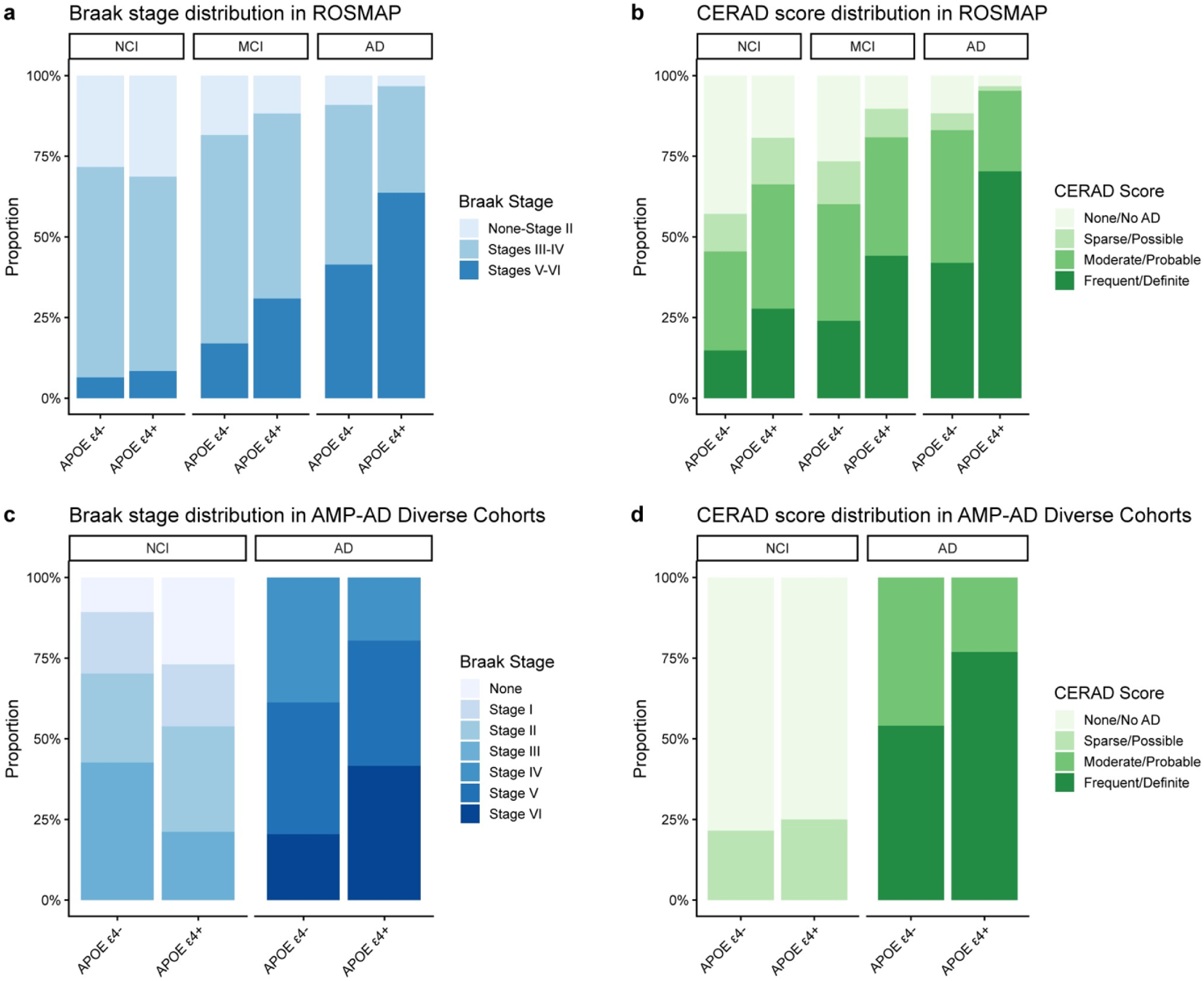
Neuropathological burden in *APOE* ε4 carriers and non-carriers across diagnostic groups. (a-b) Braak stage and CERAD score distributions in ROSMAP, respectively. (c-d) Braak stage and CERAD score distributions in the AMP-AD Diverse Cohorts Study, respectively. AD, Alzheimer’s disease; AMP-AD, Accelerating Medicines Partnership-Alzheimer’s Disease; APOE, apolipoprotein E; CERAD, Consortium to Establish a Registry for Alzheimer’s Disease; MCI, mild cognitive impairment; NCI, no cognitive impairment; ROSMAP, Religious Orders Study and Memory and Aging Project.

To assess the robustness of these findings in a more ancestrally diverse sample with neuropathologically confirmed diagnoses, we examined the same associations in the AMP-AD Diverse Cohorts Study. Consistent with the ROSMAP results, *APOE* ε4 carriage was significantly associated with higher Braak stage in the AD group (OR = 2.03, 95% CI: 1.50 - 2.77) but not in the NCI group (OR = 0.73, 95% CI: 0.41 - 1.32) (Fig. 4c). Similarly, *APOE* ε4 carriage was significantly associated with more severe CERAD scores in the AD group (OR = 2.07, 95% CI: 1.44 - 3.01) but not in the NCI group (OR = 1.86, 95% CI: 0.87 - 3.92) (Fig. 4d). Together, these findings indicate that *APOE* ε4 carriers with cognitive impairment show greater neurofibrillary tangle and neuritic plaque burden relative to non-carriers.

### *APOE* ε4-associated transcriptomic and proteomic changes in post-mortem brain tissue are minimal and region-specific

To explore the molecular basis of the neuropathological changes observed in *APOE* ε4 carriers, we examined whether *APOE* ε4 carriers exhibit distinct transcriptomic and proteomic signatures in bulk post-mortem brain tissue. Transcriptomic changes in four brain regions (dlPFC, hCN, PCC, and STG) were assessed across the ROSMAP cohort and the AMP-AD Diverse Cohorts Study using models adjusted for age at death, sex, diagnosis, and batch (Fig. 5a). There were limited *APOE* ε4-associated transcriptomic changes identified across all regions. In the ROSMAP cohort, only one DEG was detected in the dlPFC, six in the hCN, and five in the PCC (FDR < 0.05; Fig. 5b-d). Across brain regions, MTCO2P12 was upregulated, and CHI3L2 was downregulated in both the hCN and PCC, but not the dlPFC. In the AMP-AD Diverse Cohorts Study, five DEGs were identified in the dlPFC and 20 in the STG (Fig. 5e-f). Among them, only MTCO1P12 was upregulated in both regions. In addition, MI-based feature selection identified no genes exceeding a threshold of 0.1 in any region across both cohorts, further supporting the absence of a strong *APOE* ε4-associated transcriptomic signal in bulk brain tissue (Supplementary Fig. 1). *APOE* ε4-associated proteomic changes were characterised in the dlPFC of both cohorts (Fig. 5a). A linear model adjusted for age at death, sex, diagnosis, and batch identified nine *APOE* ε4-associated DAPs in the ROSMAP dlPFC (FDR < 0.05; Fig. 5g). In the AMP-AD Diverse Cohorts Study, the same analysis identified eight DAPs in the dlPFC. Across both cohorts, five of these overlapped (MDK, COL25A1, APP, HTRA1, SPOCK2), indicating robust changes independent of cohort (Fig. 5h). MI-based feature selection similarly revealed weak *APOE* ε4-associated proteomic signals across both cohorts (Supplementary Fig. 2).

**Figure 5.**
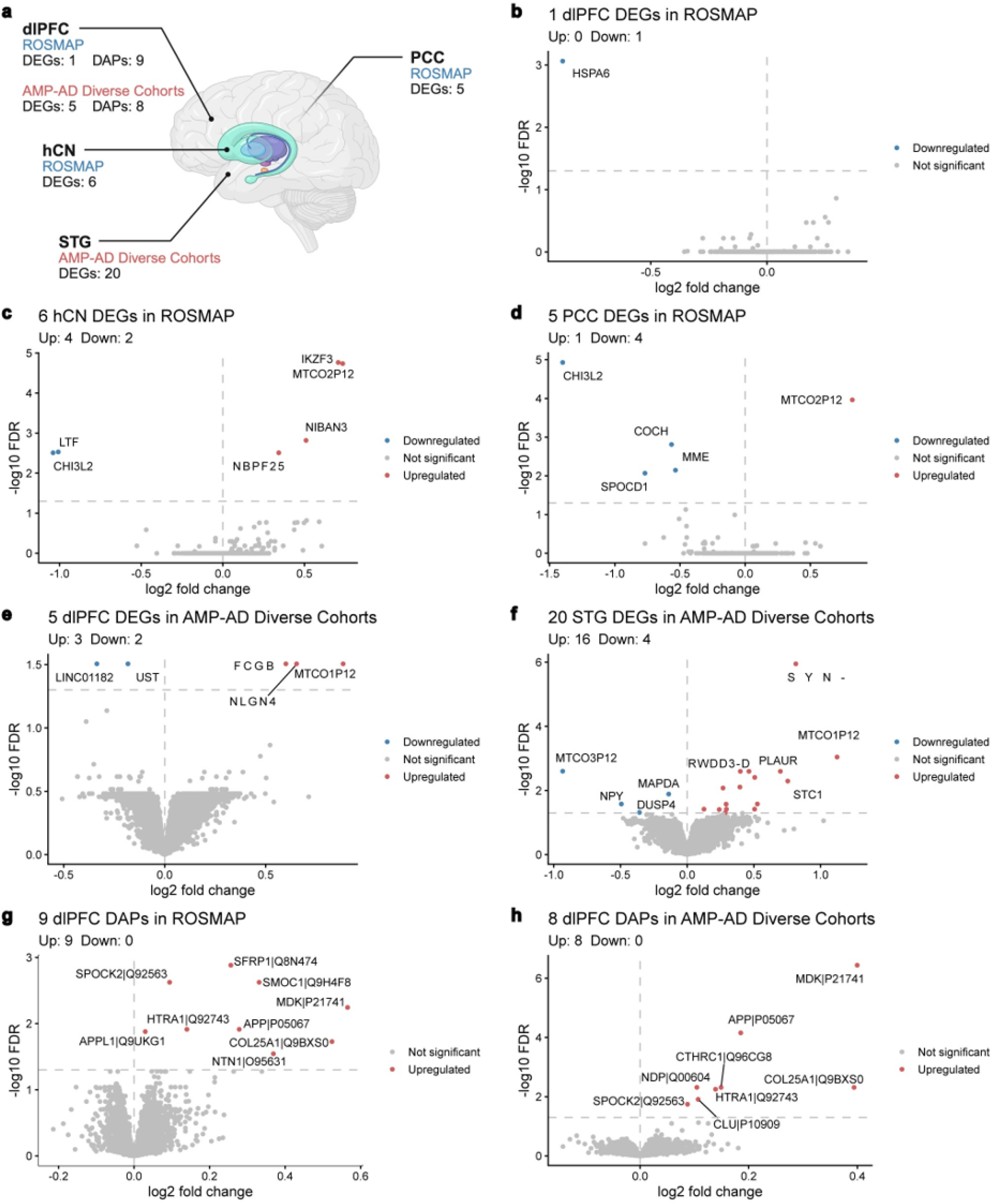
*APOE* ε4-associated transcriptomic and proteomic changes in bulk post-mortem brain tissue. (a) Overview of *APOE* ε4-associated DEGs and DAPs identified in different brain regions. (b-d) Volcano plots of *APOE* ε4-associated DEGs in the ROSMAP dlPFC, hCN, and PCC, respectively (FDR < 0.05). (e-f) Volcano plots of *APOE* ε4-associated DEGs in the AMP-AD Diverse Cohorts Study dlPFC and STG, respectively (FDR < 0.05). (g-h) Volcano plots of *APOE* ε4-associated DAPs in the ROSMAP and AMP-AD Diverse Cohorts Study dlPFC, respectively (FDR < 0.05). AMP-AD, Accelerating Medicines Partnership-Alzheimer’s Disease; APOE, apolipoprotein E; DAP, differentially abundant protein; DEG, differentially expressed gene; dlPFC, dorsolateral prefrontal cortex; FDR, false discovery rate; hCN, head of the caudate nucleus; PCC, posterior cingulate cortex; ROSMAP, Religious Orders Study and Memory and Aging Project; STG, superior temporal gyrus.

Together, these findings suggest that *APOE* ε4-associated transcriptomic changes in bulk post-mortem brain tissue are minimal and largely region-specific, with only a small number of DEGs shared across brain regions. *APOE* ε4-associated proteomic changes in the dlPFC were similarly modest but reproducible across cohorts.

### Limited concordance between transcriptomic and proteomic changes in brain tissue is not specific to *APOE* ε4

We next sought to investigate whether *APOE* ε4-associated mRNA changes in brain tissue are reflected at the protein level. To do this, we identified 466 AMP-AD Diverse Cohorts Study participants with paired transcriptomic and proteomic data for the dlPFC. Among mRNAs and proteins mapped to the same gene symbol, Spearman correlation of log2 fold changes revealed weak-to-moderate correlation (r = 0.455, *p* < 0.001) between *APOE* ε4-associated mRNA and protein changes (Fig. 6a). To determine whether this pattern was specific to *APOE* ε4, we repeated the analysis using AD-associated log2 fold changes and found a comparable weak-to-moderate correlation (r = 0.329, *p* < 0.001) (Fig. 6b). Together, these results suggest that limited mRNA-protein concordance is likely a general feature of bulk brain tissue, rather than a specific consequence of *APOE* ε4 carriage.

**Figure 6.**
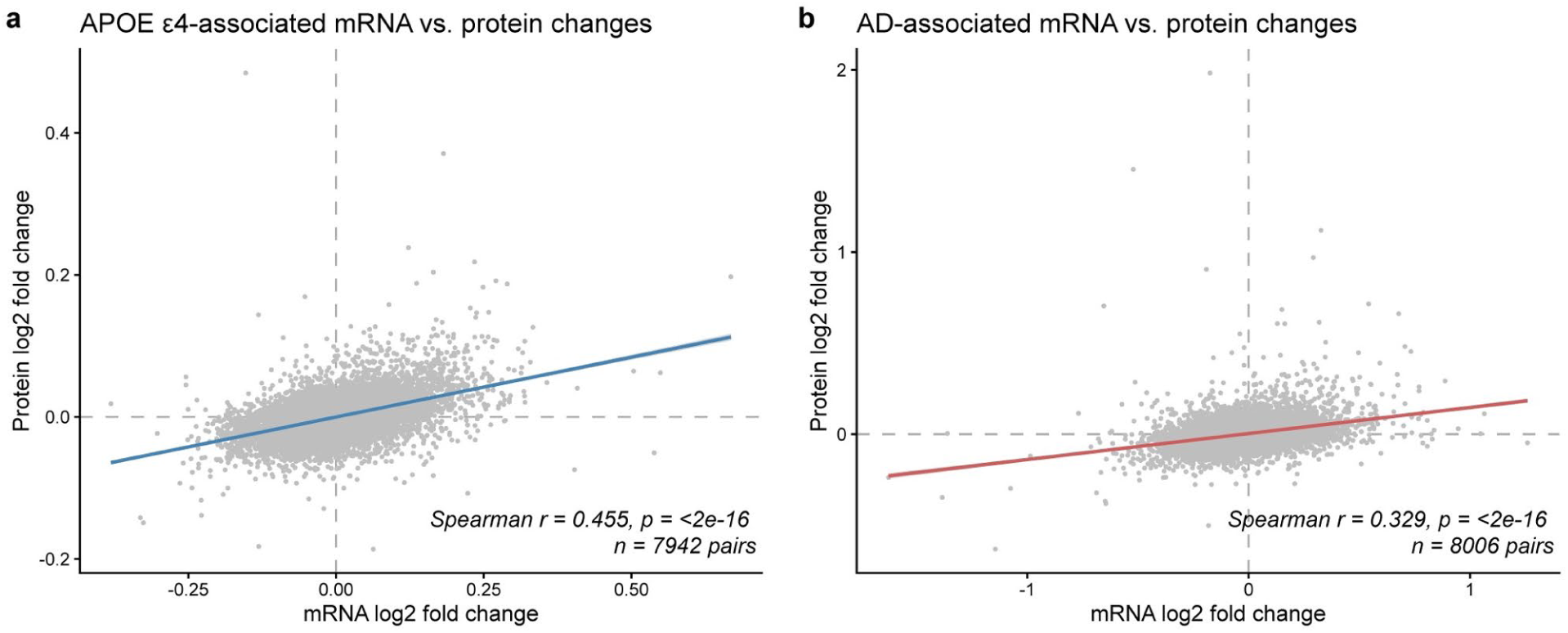
mRNA-protein concordance of *APOE* ε4-and AD-associated changes in the dlPFC of the AMP-AD Diverse Cohorts Study. Scatter plots of log_2_ fold changes between matched transcriptomic and proteomic data for (a) *APOE* ε4-and (b) AD-associated changes in the dlPFC. AD, Alzheimer’s disease; AMP-AD, Accelerating Medicines Partnership-Alzheimer’s Disease; APOE, apolipoprotein E; dlPFC, dorsolateral prefrontal cortex.

### *APOE* ε4-associated proteomic signatures are detectable in iPSC-derived astrocytes and neurons

Given the limited *APOE* ε4-associated proteomic signal observed in post-mortem brain tissue, we next sought to determine whether this reflects a genuine absence of effect or a limitation of examining bulk tissue. To do so, we analysed proteomic data from iAstrocytes and iNeurons generated from ROSMAP participants to obtain a cell type-specific view of *APOE* ε4-associated proteomic changes. Linear models adjusted for diagnosis and sex identified no significant DAPs in either iAstrocytes or iNeurons (Supplementary Fig. 3). In contrast, MI-based feature selection identified 37 proteins with non-zero association with *APOE* ε4 carriage in iAstrocytes and 43 in iNeurons (Fig. 7a-b). PCA based on the top 13 MI-selected proteins for each cell type revealed clear separation by *APOE* ε4 carriage in both datasets, indicating that *APOE* ε4-associated proteomic changes are more readily detectable at the cellular level than in bulk tissue (Fig. 7c-d). Notably, no MI-selected proteins were shared between the two cell types, suggesting that *APOE* ε4-associated proteomic changes are strongly cell type-specific.

**Figure 7.**
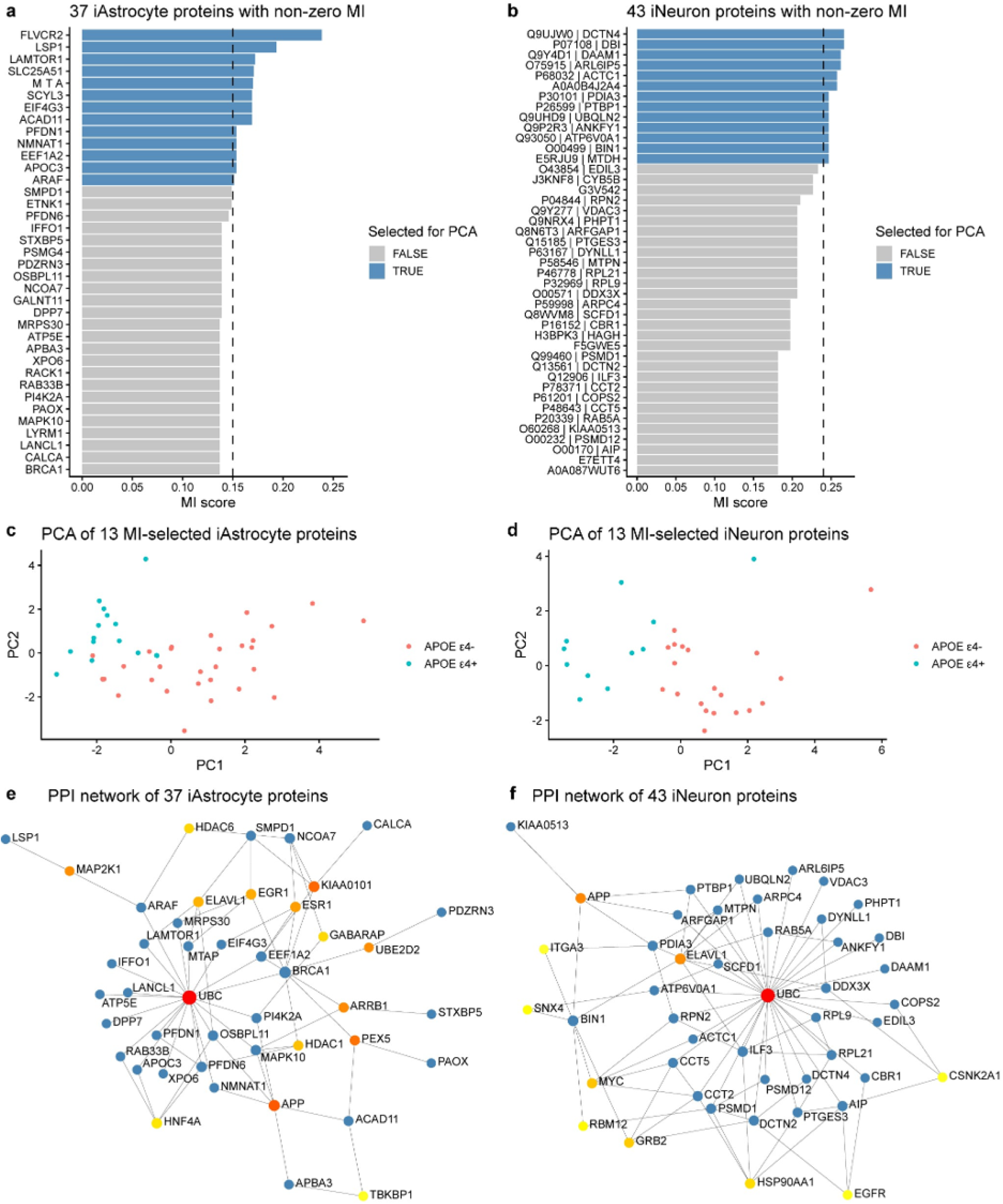
*APOE* ε4-associated proteomic signatures in ROSMAP iPSC-derived astrocytes and neurons. (a-b) Bar plots of MI scores for 37 iAstrocyte and 43 iNeuron proteins with non-zero MI for *APOE* ε4 carriage, respectively. (c-d) PCA of the top 13 MI-selected iAstrocyte and iNeuron proteins by *APOE* ε4 carriage, respectively. (e-f) Protein-protein interaction networks of the 37 *APOE* ε4-associated iAstrocyte and 43 *APOE* ε4-associated iNeuron proteins, respectively. APOE, apolipoprotein E; iPSC, induced pluripotent stem cell; MI, mutual information; PCA, principal component analysis; ROSMAP, Religious Orders Study and Memory and Aging Project.

Protein-protein interaction network analysis of the 37 *APOE* ε4-associated iAstrocyte proteins and 43 iNeuron proteins revealed strong functional connectivity in both cell types (Fig. 7e-f). However, no significant enrichment of GO biological processes or Reactome pathways was identified in either dataset after FDR correction (Supplementary Tables 8-9).

Taken together, these findings suggest that *APOE* ε4-associated proteomic signatures are present in both iAstrocytes and iNeurons, and that the limited signal observed in bulk post-mortem brain tissue is likely attributable to the dilution of cell type-specific effects within heterogeneous tissue.

## Discussion

In this study, we characterised *APOE* ε4-associated transcriptomic and proteomic changes across plasma, CSF, post-mortem brain tissue, and iPSC-derived astrocytes and neurons. In plasma, we identified an immune-enriched proteomic signature in *APOE* ε4 carriers. This signature was independent of AD diagnosis and generalised to CSF in an independent cohort. Mediation analysis further revealed a subset of *APOE* ε4-associated plasma proteins that may mediate or reflect downstream consequences of AD pathology. Most notably, CTF1 appeared to mediate the relationship between *APOE* ε4 and AD, with both *APOE* ε4 carriage and AD pathology associated with increased CTF1 abundance. In contrast, the *APOE* ε4-associated reduction in NEFL appeared to be partially offset by AD-driven upregulation. In the brain, *APOE* ε4 carriage was associated with greater neuritic plaque burden across all diagnostic groups in ROSMAP, and this association was replicated in individuals with AD in the AMP-AD Diverse Cohorts. *APOE* ε4 carriers also exhibited greater neurofibrillary tangle burden, but only in individuals with cognitive impairment in both cohorts. In contrast, relatively few *APOE* ε4-associated transcriptomic and proteomic changes were detected in bulk brain tissue, with poor concordance between the two molecular layers. This likely reflects the dilution of cell type-specific signals in bulk tissue rather than a true absence of biological effect, as supported by distinct *APOE* ε4-associated proteomic signatures detected in iPSC-derived astrocytes and neurons. Together, these findings indicate that *APOE* ε4 is associated with a consistent proteomic signature observed across biofluids. Its effects in the brain are more complex and vary across cell types and molecular layers.

A key finding of our study is that *APOE* ε4-associated proteomic changes are shared across plasma and CSF. A recent study by Seo et al. reported a moderate correlation between *APOE* ε4-associated protein changes across these biofluids (17). We extend this by using machine learning to demonstrate that the top *APOE* ε4-associated proteins identified in plasma, including SPC25 and S100A13, can reliably discriminate *APOE* ε4 carriers from non-carriers when measured in CSF. This cross-biofluid generalisability supports previous pathway-level analyses showing convergent dysregulation of immune, gene regulatory, and metabolic processes in both biofluids (6, 9). Notably, the plasma and CSF measurements were obtained from two independent cohorts, providing external validation and evidence that these findings are not cohort specific. In addition, we confirmed that this proteomic signature was independent of both sex and disease stage, consistent with previous studies (8, 9, 51). Together, our findings suggest that *APOE* ε4 confers a systemic, AD-independent proteomic signature across biofluids, closely associated with immune dysregulation.

Furthermore, using mediation analysis, we identified a subset of *APOE* ε4-associated plasma proteins that may bidirectionally interact with AD. Among these, CTF1 was significantly upregulated in *APOE* ε4 carriers, consistent with previous reports (6, 51). This upregulation of CTF1 was statistically associated with the relationship between *APOE* ε4 and increased AD risk. Our analysis further revealed that AD status may also mediate the effect of *APOE* ε4 on CTF1 abundance, consistent with prior reports of elevated CTF1 levels in individuals with AD (51, 52). Together, these findings suggest with a bidirectional statistical association in which *APOE* ε4-associated upregulation of CTF1 is associated with AD risk, while AD pathology is also associated with increased CTF1 abundance in *APOE* ε4 carriers. In contrast, NEFL was significantly downregulated in *APOE* ε4 carriers and mediated a reduction in AD risk. Previous studies have reported inconsistent effects of *APOE* ε4 on plasma NEFL levels, with some observing a decrease (6, 9), others an increase (53, 54), and others no significant difference (55). Our findings provide one possible explanation for this discrepancy. While *APOE* ε4 is associated with a reduction in NEFL, AD pathology may mediate an increase in NEFL abundance, a pattern consistent with the well-established elevation of NEFL in AD reflecting neuronal injury (56, 57, 58). Depending on a cohort’s genotype and disease stage composition, the AD-associated increase in NEFL levels in *APOE* ε4 carriers may therefore mask or offset the *APOE* ε4-associated reduction. Together, this highlights the complex relationship between *APOE* ε4 and AD and illustrates how genotype and disease status may jointly shape plasma proteomic signals.

Consistent with earlier studies linking *APOE* ε4 to increased amyloid and tau pathology (18, 19), we observed greater neuritic plaque and neurofibrillary tangle burden among *APOE* ε4 carriers in both cohorts. In ROSMAP, *APOE* ε4 carriage was associated with more severe neuritic plaque burden across all diagnostic groups, whereas neurofibrillary tangle burden was elevated only among individuals with MCI or AD. This pattern is consistent with evidence that amyloid accumulation may precede and contribute to subsequent tau pathology in *APOE* ε4 carriers (59). In the AMP-AD Diverse Cohorts Study, however, increased neuritic plaque and neurofibrillary tangle burden were observed only among *APOE* ε4 carriers with AD. This divergence may reflect differences in disease-stage classification and cohort composition. Specifically, the absence of an intermediate MCI category in the AMP-AD Diverse Cohorts Study could limit the detection of pathological differences at earlier disease stages. The higher proportion of Black and African American participants in the AMP-AD Diverse Cohorts Study relative to the predominantly White ROSMAP cohort may also contribute to this divergence. This is consistent with a previous study showing the strongest *APOE* ε4-associated effects on AD-related burden in individuals of European ancestry (60).

In contrast to the clear association between *APOE* ε4 and AD pathology, we identified relatively few *APOE* ε4-associated transcriptomic and proteomic changes in bulk post-mortem brain tissue. This likely reflects the dilution of cell type-specific signals in heterogeneous bulk tissue. Previous studies using single-cell RNA-seq have identified *APOE* ε4-associated transcriptomic changes in the brain that are largely cell type-specific (11, 12, 15). In contrast, bulk RNA-seq analyses have identified comparatively fewer changes (19). Our proteomic analyses of iPSC-derived astrocytes and neurons further support this cell type-specific view by revealing distinct *APOE* ε4-associated signatures in each cell type. This is consistent with previous studies reporting that *APOE* ε4-associated changes are more pronounced in iAstrocytes, implicating mitochondrial and inflammatory functions (61, 62, 63), and comparatively less pronounced in neurons, where they implicate inflammation and neuronal signalling (63). To our knowledge, this is also the first study to directly compare *APOE* ε4-associated transcriptomic changes across multiple human brain regions. The marked region specificity we observe may reflect differences in cellular composition across regions or regional variation in the effect of *APOE* ε4 itself and warrants investigation in future studies.

Previous studies have highlighted limited concordance between mRNA and protein abundance in the brain, indicating that transcriptomic changes are not reliable proxies for downstream protein-level effects (20, 21, 22). Our study extends this by showing that correlations between mRNA and protein changes associated with both AD and *APOE* ε4 are similarly moderate, ranging from 0.3 to 0.5. A previous study examining AD-associated mRNA-protein concordance reported a comparable correlation of 0.38 (64). Johnson et al. further showcased that many AD-associated proteomic co-expression modules were absent at the transcript level (23). To our knowledge, ours is the first study to examine mRNA-protein concordance in relation to *APOE* ε4. Together, these findings highlight that transcriptomic profiling alone is insufficient to capture genotype or disease-associated molecular mechanisms in brain tissue. Future studies integrating proteomic and other molecular profiling will provide a more comprehensive biological insight.

Despite these insights, several limitations warrant consideration. First, although sample sizes were large across most analyses, homozygous *APOE* ε4 carriers remain relatively rare, particularly among cognitively unimpaired individuals. While this rarity reflects the strong link between homozygous carriage and AD risk, it limited our ability to examine dose-dependent effects of *APOE* ε4. Future studies incorporating cohorts enriched for homozygous carriers would help address this. Second, our mediation analyses were conducted on cross-sectional observations. While they offer insight into mediation relationships between *APOE* ε4, plasma proteins, and AD status, these should be interpreted as statistical associations rather than causal inferences. Longitudinal studies with repeated protein measurements would help clarify the true temporal sequence of these relationships.

## Conclusions

In conclusion, our study provides a comprehensive investigation of *APOE* ε4-associated molecular changes across biofluids, brain tissue, and iPSC-derived cells. We showed that *APOE* ε4 is associated with a systemic, immune-related proteomic signature that is consistent across plasma and CSF, and independent of AD diagnosis and sex. This raises the possibility that plasma could serve as a more accessible alternative to CSF for studying *APOE* ε4-related biology, with potential implications for risk stratification or therapeutic targeting in *APOE* ε4 carriers. In the brain, *APOE* ε4-associated transcriptomic and proteomic signatures appear to be largely cell type-specific, with limited concordance observed between the two molecular layers. Our findings therefore underscore the value of integrating multi-omic and cell type-resolved approaches to advance understanding of the molecular mechanisms linking *APOE* ε4 to AD pathogenesis.

## List of abbreviations

ACME: average causal mediation effect
AD: Alzheimer’s disease
ADNI: Alzheimer’s Disease Neuroimaging Initiative
AMP-AD: Accelerating Medicines Partnership – Alzheimer’s Disease
ANML: adaptive normalization by maximum likelihood
APOE: apolipoprotein E
AUC: area under the curve
CDR: Clinical Dementia Rating
CERAD: Consortium to Establish a Registry for Alzheimer’s Disease
CI: confidence interval
CSF: cerebrospinal fluid
DAP: differentially abundant protein
DEG: differentially expressed gene
FDR: false discovery rate
hCN: head of caudate nucleus
IMEx: International Molecular Exchange Consortium
dlPFC: dorsolateral prefrontal cortex
GO: Gene Ontology
iPSC: induced pluripotent stem cell
MCI: mild cognitive impairment
MI: mutual information
MMSE: Mini-Mental State Exam
NCI: no cognitive impairment
NPV: negative predictive value
OR: odds ratio
PBMCs: peripheral blood mononuclear cells
PCA: principal component analysis
PCC: posterior cingulate cortex
PPV: positive predictive value
RFU: relative fluorescent units
ROSMAP: Religious Orders Study and Rush Memory and Aging Project
SMOTE: Synthetic Minority Oversampling Technique
STG: superior temporal gyrus

## Declarations

### Ethics approval and consent to participate

The included studies were approved by their respective Institutional Review Boards (IRB) and participants provided informed written consent to participate.

### Consent for publication

N/A

### Availability of data and materials

The ROSMAP and AMP-AD Diverse Cohorts Study data are available through the AD Knowledge Portal (https://adknowledgeportal.synapse.org/). Researchers who wish to access these controlled datasets are required to submit a Data Use Agreement. More information can be found here: https://adknowledgeportal.synapse.org/Data%20Access. The ADNI data are available upon request through the ADNI database (https://ida.loni.usc.edu/). Details on how to request access are available at https://adni.loni.usc.edu/data-samples/adni-data/#AccessData. All code used in this study is publicly available at https://github.com/Christy-lll/APOE_AD.

### Competing interests

The authors declare no competing interests.

### Funding

This work was supported by the Australian Government’s Medical Research Future Fund MRF2040081 (C.A.F. & A.S.), MRF2052401 (A.S. & C.A.F.); and philanthropic funding from John & Anne Leece Family (A.S.), Paul & Valeria Ainsworth Family (C.A.F.), and Neil and Norma Hill Foundation (C.A.F).

### Authors’ contributions

Conceptualization, data curation, funding acquisition, project administration, resources, supervision, validation, writing – review & editing: C.A.F. and A.S. Formal analysis, investigation, methodology, software, visualization: X.C.L. and A.S. Writing – original draft: X.C.L.

## Acknowledgements

The results published here are in whole or in part based on data obtained from The AD Knowledge Portal (https://doi.org/10.7303/9618239). Study data were provided by the Rush Alzheimer’s Disease Center, Rush University Medical Center, Chicago. Data collection was supported through funding by NIA grants P30AG10161 (ROS), R01AG15819 (ROSMAP; genomics and RNAseq), R01AG17917 (MAP), R01AG30146, R01AG36042 (5hC methylation, ATACseq), RC2AG036547 (H3K9Ac), R01AG36836 (RNAseq), R01AG48015 (monocyte RNAseq) RF1AG57473 (single nucleus RNAseq), U01AG32984 (genomic and whole exome sequencing), U01AG46152 (ROSMAP AMP-AD, targeted proteomics), U01AG46161(TMT proteomics), U01AG61356 (whole genome sequencing, targeted proteomics, ROSMAP AMP-AD), P30AG072975, the Illinois Department of Public Health (ROSMAP), and the Translational Genomics Research Institute (genomic). Additional phenotypic data can be requested at www.radc.rush.edu. Study data were provided through the Accelerating Medicine Partnership for AD (U01AG046161 and U01AG061357) based on samples provided by the Rush Alzheimer’s Disease Center, Rush University Medical Center, Chicago.

Data collection was supported through funding by NIA grants P30AG10161, R01AG15819, R01AG17917, R01AG30146, R01AG36836, U01AG32984, U01AG46152, the Illinois Department of Public Health, and the Translational Genomics Research Institute.

Annie J. Lee, Yiyi Ma, Lei Yu, Robert J. Dawe, Cristin McCabe, Konstantinos Arfanakis, Richard Mayeux, David A. Bennett, Hans-Ulrich Klein, and Philip L. De Jager. Multi-region brain transcriptomes uncover two subtypes of aging individuals with differences in Alzheimer’s disease risk and the impact of APOEε4. bioRxiv 2021.

The results published here are in whole or in part based on data obtained from the AD Knowledge Portal Diverse Cohort Study DOI (https://doi.org/10.7303/9618093). Data generation was supported by the following NIH grants: U01AG046139, U01AG046170, U01AG061357, U01AG061356, U01AG061359, and R01AG067025. We thank the participants of participants of the Religious Order Study, Memory and Aging Project, the Minority Aging Research Study, Rush Alzheimer’s Disease Research Center, Mount Sinai/JJ Peters VA Medical Center NIH Brain and Tissue Repository, National Institute of Mental Health Human Brain Collection Core (NIMH HBCC), Mayo Clinic Brain Bank, Sun Health Research Institute Brain and Body Donation Program, Goizueta Alzheimer’s Disease Research Center, New York Brain Bank at Columbia University, New York Genome Center and the Biggs Institute Brain Bank for their generous donations. Data and analysis contributing investigators include Nilüfer Ertekin-Taner, Minerva Carrasquillo, Mariet Allen (Mayo Clinic, Jacksonville, FL), David Bennett, Lisa Barnes (Rush University), Philip De Jager, Vilas Menon (Columbia University), Bin Zhang, Vahram Haroutanian (Icahn School of Medicine at Mount Sinai), Allan Levey, Nick Seyfried (Emory University), Rima Kaddurah-Daouk (Duke University), Steve Finkbeiner (University of California-San Francisco/Gladstone Institutes), Daifeng Wang (University of Wisconsin-Madison), Stefano Marenco (NIMH HBCC), Anna Greenwood, Abby Vander Linden, Laura Heath, William Poehlman (Sage Bionetworks).

## Notes

### Competing Interest Statement

The authors have declared no competing interest.

