## Supplementary Figures for "Proteomic signatures of *APOE* ε4 across human tissues and cell types in Alzheimer’s disease"

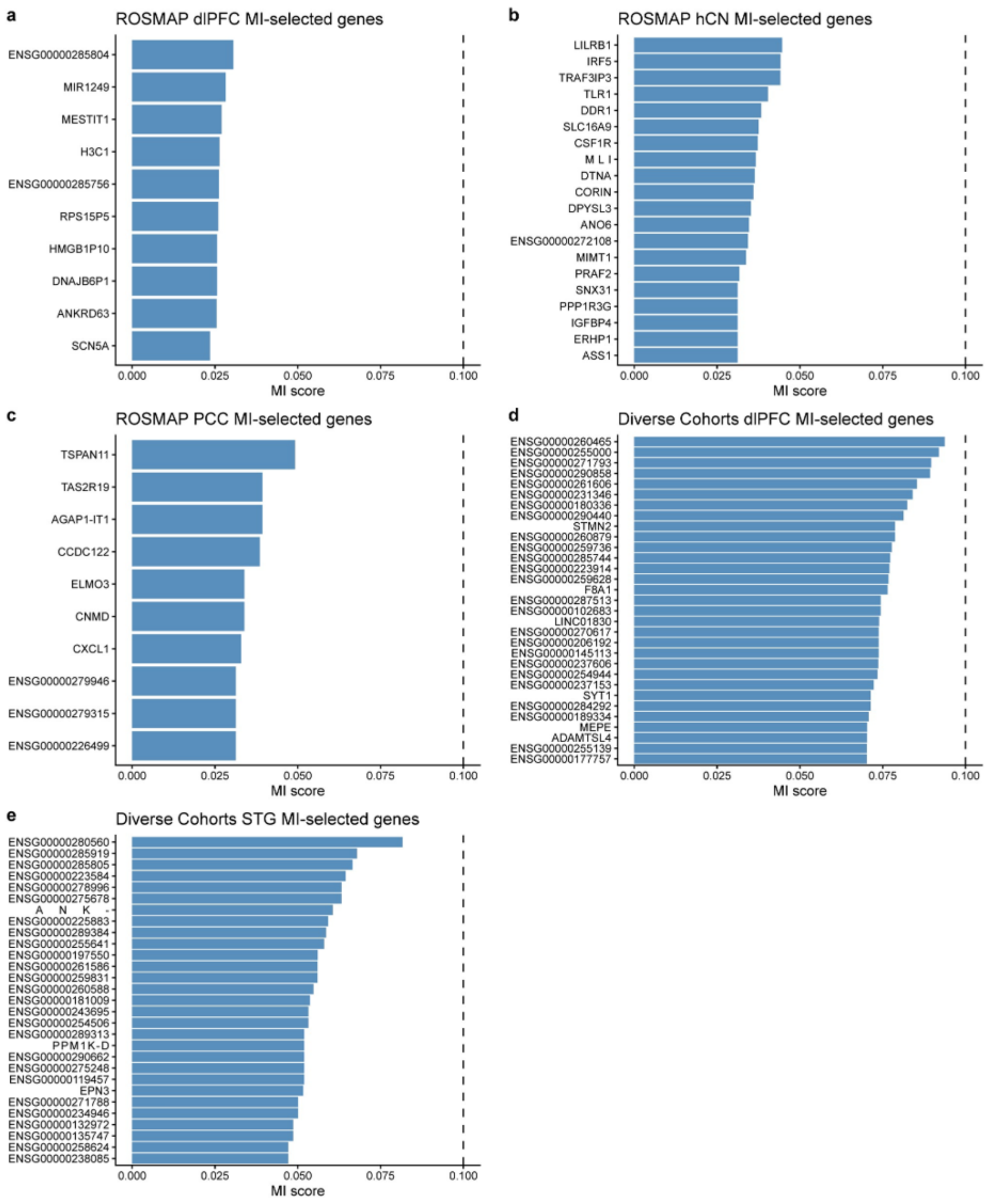


**Supplementary Figure 1. Mutual information analysis of *APOE* ε4-associated transcriptomic changes in bulk post-mortem brain tissue.** Bar plots show genes with non-zero MI scores for *APOE* ε4 carriage; where more than 30 genes had non-zero scores, only the top 30 are displayed. The vertical line indicates an MI score of 0.1. (a-c) MI scores in the ROSMAP dlPFC, hCN, and PCC, respectively. (d-e) MI scores in the AMP-AD Diverse Cohorts Study dlPFC and STG, respectively. AMP-AD, Accelerating Medicines Partnership - Alzheimer's Disease; APOE, apolipoprotein E; dlPFC, dorsolateral prefrontal cortex; hCN, head of the caudate nucleus; MI, mutual information; PCC, posterior cingulate cortex; ROSMAP, Religious Orders Study and Memory and Aging Project; STG, superior temporal gyrus.

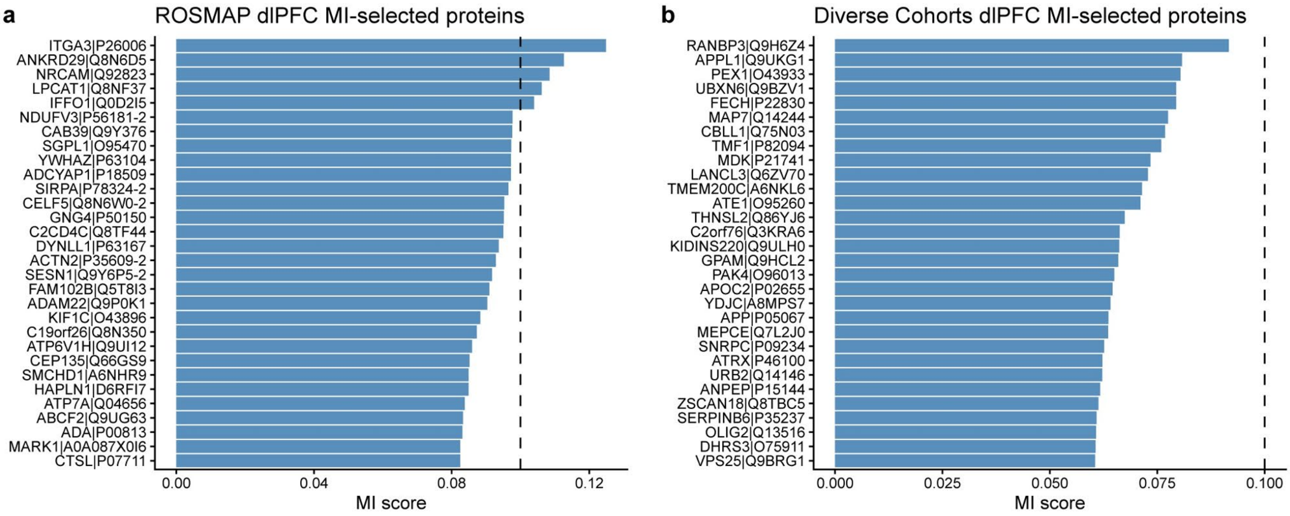


**Supplementary Figure 2. Mutual information analysis of *APOE* ε4-associated proteomic changes in the dlPFC.** Bar plots show proteins with non-zero MI scores for *APOE* ε4 carriage; where more than 30 genes had non-zero scores, only the top 30 are displayed. The vertical line indicates an MI score of 0.1. (a) MI scores in the ROSMAP dlPFC. (b) MI scores in the AMP-AD Diverse Cohorts Study dlPFC. AMP-AD, Accelerating Medicines Partnership - Alzheimer's Disease; APOE, apolipoprotein E; dlPFC, dorsolateral prefrontal cortex; MI, mutual information; ROSMAP, Religious Orders Study and Memory and Aging Project.

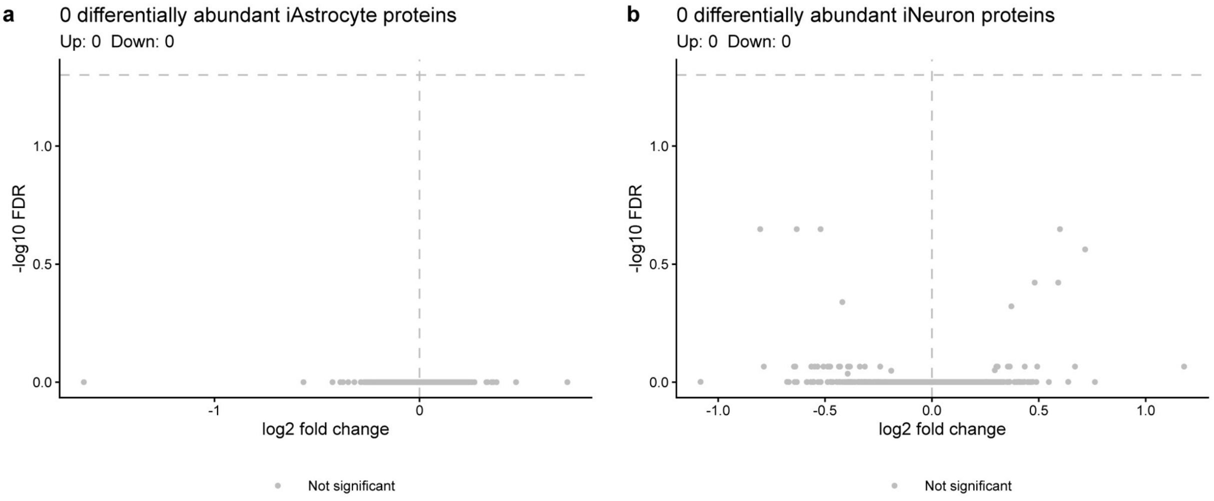


**Supplementary Figure 3. Differential abundance analysis of *APOE* ε4-associated proteomic changes in ROSMAP iAstrocytes and iNeurons.** Volcano plots show proteins tested for differential abundance by *APOE* ε4 carriage. No significant DAPs were identified in either (a) iAstrocytes or (b) iNeurons (FDR < 0.05). APOE, apolipoprotein E; DAP, differentially abundant protein; FDR, false discovery rate; ROSMAP, Religious Orders Study and Memory and Aging Project.
